# Epromoter 3D interaction-associated regulation in T-acute Lymphoblastic Leukemia

**DOI:** 10.64898/2026.08.25.746923

**Authors:** Joaquin Felipe Roca Paixão, Iris Manosalva, Antoine Pinton, Agata Cieslak, Carla Cardone, Nathalie Sakakini, Nori Sadouni, Andreas Zanzoni, Guillaume Andrieu, Vahid Asnafi, Aurore Touzart, Salvatore Spicuglia

**Affiliations:** Aix-Marseille University, INSERM, TAGC, UMR 1090, Marseille, France; Université Paris Cité, Institut Necker-Enfants Malades (INEM), Institut national de la santé et de la recherche médicale (Inserm) U1151), Paris, France; Laboratory of Onco-Hematology, Hôpital Necker, Assistance Publique-Hôpitaux de Paris (AP-HP), Paris, France

**Keywords:** Epromoter, promoter-promoter interactions, HiChIP, T-ALL, 3D interactions

## Abstract

**Background:** Promoters have been traditionally seen as contiguous gene-adjacent cis-regulatory elements. Yet, substantial studies corroborate that Epromoters (promoters with enhancer activity) engage in distal forms of gene regulation. Although in the three-dimensional (3D) space enhancer-promoter networks have been well studied, the contribution of the circuits of promoter-promoter (P-P) interactions is poorly understood. Furthermore, whether the regulatory aspects of P-P interactions in cancer may be controlled by physical 3D-mediated Epromoter interactions remains elusive.

**Results:** We show that Epromoter-mediated 3D interactions regulate target genes and participate in cluster co-regulation, playing a critical role in T-cell acute Lymphoblastic Leukemia (T-ALL). To achieve this, we first leveraged survival CRISPR screenings in T-ALL model cells (Jurkat) to identify potential Epromoters. By integrating these findings with an H3K27ac HiChIP dataset from T-ALL cells, we characterized a set of Epromoters that establish 3D genome interactions with other promoters. We observed that promoters organize into dense, promoter-rich genomic clusters, and that among them, the clusters enriched with Epromoters actively regulate complex gene expression networks. To investigate gene coregulation, we integrated transcriptomic data from T-ALL patients and found that promoter-promoter (P-P) pairs exhibit positive correlation at multiple levels, and that several Jurkat Epromoter candidate clusters are significantly co-regulated in the patient cohort. To experimentally validate these candidates, we utilized CRISPRi to inhibit Epromoters, which revealed direct transcriptional regulation of multiple target genes within each hub. Finally, we performed cell competition assays to confirm that these Epromoters are vital for T-ALL cell survival.

**Conclusions:** Our analysis provides support for the role of Epromoters in the regulation of 3D P-P interactions and co-regulation of promoter hubs, and how these interactions play a critical part in T-ALL cell survival.

## Background

The three-dimensional (3D) architecture of chromatin forms a highly structured organization with distinct layers of compartmentalization in the nuclear space, depicting developmental and tissue-specific cell states [1–3]. Techniques used to untangle 3D genome organization collectively known as chromosome conformation capture have undergone rapid methodological development and diversification, with several approaches specifically optimized to capture interactions between regulatory elements, such as Capture Hi-C, ChIAP-PET and H3K27ac HiChIP [4,5]. At the megabase scale, chromatin folds into topologically associating domains (TADs), where intra-region proximity is greater than inter-region, driving long-range interactions within the TAD, in which clusters of cis-regulatory elements frequently form interaction hubs [6–8]. One of the key aspects is that distal enhancers (E) and promoters (P) form specific physical interactions whose strength and range are tuned in a cell-state-dependent manner [9–11] and subtle changes in these structures may create large changes in transcription programs [12]. Clear evidence that E-P loops vary substantially across cancers has now been provided using H3K27ac HiCHIP and related 3D genomics. For instance, Yost et al. mapped cancer-type-specific E-P connectomes across 15 primary tumor types, while Breves et al. identified hyperconnected 3D regulatory hubs in glioblastoma stem cells [13,14]. A striking example is that MYC-centered hubs exhibit cancer-specific interaction patterns with distinct sets of E and P partners [14]. Current models still emphasize enhancer activity as the principal driver of spatial gene regulation [15–17]. Yet, P-P interactions, although repeatedly observed in 3D studies, have often been underappreciated, in part due to the assumption that they mainly govern housekeeping genes [13,14,18–23].

In recent years, Epromoters have emerged as pivotal cis-regulatory sequences that behave as canonical promoters while also regulating distal genes [24–32]. Epromoters can serve as regulatory hubs that recruit stress-related transcription factors to coordinate the activation of nearby genes in response to environmental cues [33]. Moreover, they are enriched for disease-associated variants, highlighting their pleiotropic impact on multiple traits [34]. In the 3D space, promoter-centered capture assays have shown that P-P associations are enriched for expression Quantitative Trait Loci (eQTLs) and promoter capture Hi-C and H3K27ac HiChIP have demonstrated that promoter interactions can link non-coding variants to distal target genes [35,36]. These observations underscore a compelling need to investigate how Epromoters participate in P-P networks and how such interactions contribute to genetic disease.

Gene expression programs determine key aspects of cancer biology, helping to identify therapeutic biomarkers, predict patient outcomes, and characterize tumor states [37]. T-acute lymphoblastic Leukemia (T-ALL) is a genomically complex disease, driven by several genetic drivers and its different subtypes have specific gene expression profiles [38,39]. Therefore, elucidating how T-ALL genomic alterations disrupt the 3D genome architecture, particularly through the P-P alterations that have incidence in these gene expression programs, is critical to advance towards effective and precise disease-management strategies [39,40].

In this study, we ask whether Epromoter-mediated 3D interactions regulate target genes and how they contribute to T-ALL pathogenesis. First, we leveraged survival-focused CRISPR screens in the T-ALL model cell line Jurkat to identify potential Epromoters. By integrating these findings with an H3K27ac HiChIP dataset from T-ALL cells, we defined a set of putative Epromoters that establish 3D genome interactions with other promoters. We showed that promoters assemble into dense, promoter-rich genomic hubs and that hubs enriched in Epromoters actively regulate complex gene expression networks. Integrating transcriptomic data from a T-ALL patient cohort, we found that P-P pairs show significant expression correlation and that promoter hubs are significantly co-regulated in patients, with a subset of Epromoter hubs exhibiting stronger co-regulation than expected by chance. Using CRISPR interference (CRISPRi), we then inhibited selected Epromoters and demonstrated direct transcriptional control of multiple target genes within individual hubs. Finally, cell competition assays revealed that these Epromoters are essential for T-ALL cell survival, pointing to Epromoter-centered P-P hubs as key nodes whose disruption compromises leukemic cell survival.

## Results

### A CRISPR screen strategy to find essential Epromoters

We hypothesized that a set of Epromoter-like elements potentially regulating distal genes could be identified by a survival CRISPR inhibition (CRISPRi) screen, which detects essential promoters, to which we would remove the canonical essential gene promoters, determined by a CRISPR knock-out (CRISPRko) screen (Fig. 1A). We therefore performed paralleled whole-genome survival CRISPRi and CRISPRko screens using Jurkat cells stably expressing the dCas9-KRAB-MEcP2 (dCas9-KB) or the Cas9 cassettes, respectively (Fig. 1B). Upon library sequencing at days 0 and 21, we applied a threshold of −1 log_2_(fold change), corresponding to the DepMap median of common essential genes [41] and an FDR lower than 1% (Fig. 1C; Additional file 1: Tables S1a, S1b, S1c) to identify significant hits. We obtained 977 CRISPRi hits, of which 230 were also CRISPRko hits, yielding a set of 747 Epromoter-like elements (Fig. 1D). This represents 3.9% of all human promoters contained in the CRISPRi library, consistent with the average proportion of Epromoters per cell line that we had previously reported [34].

**Figure 1:**
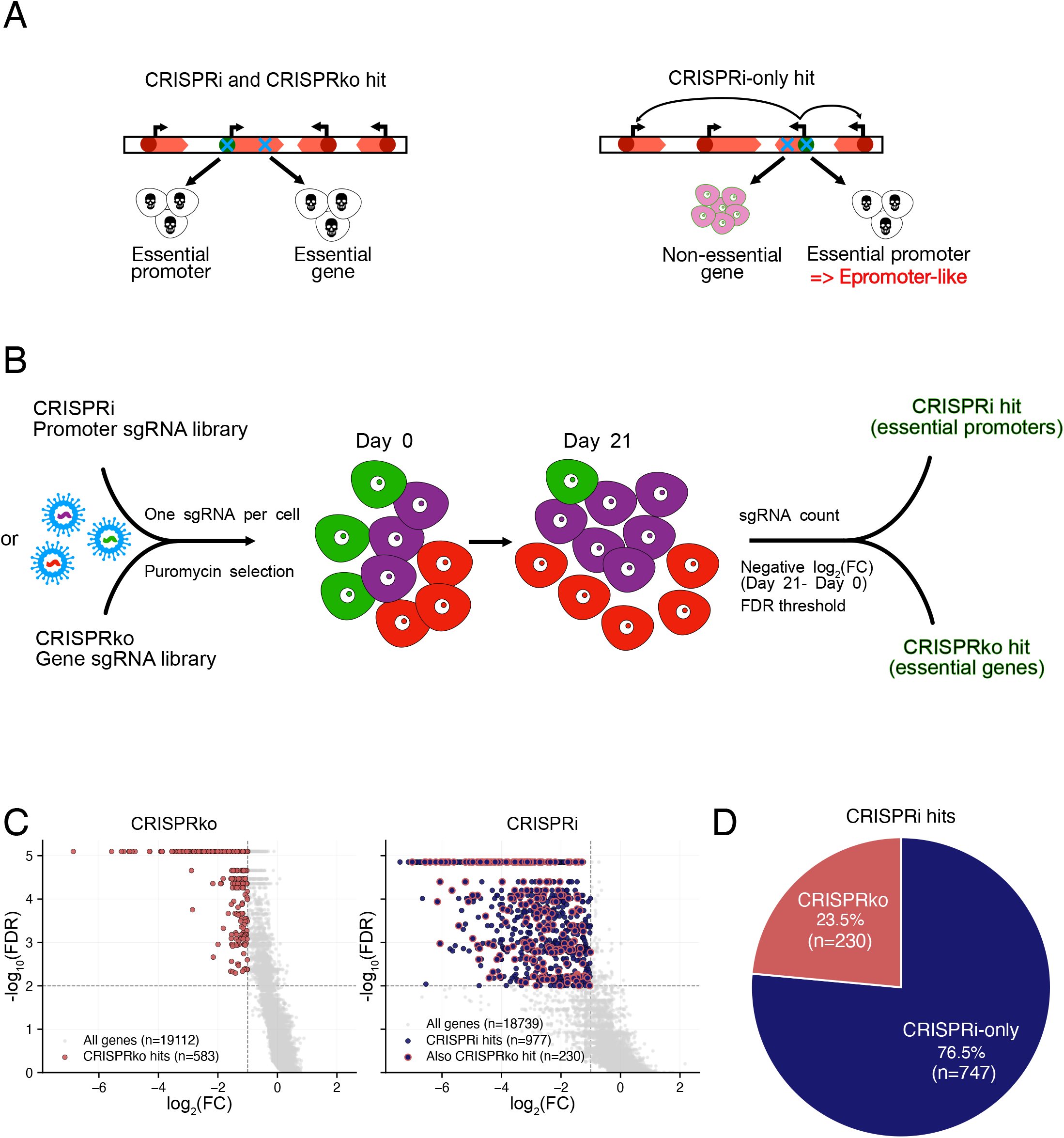
CRISPR screens identify a set of Epromoter-like elements. **A**. Schematics of the CRISPR screening strategy. Promoters identified as essential in a CRISPRi screen whose corresponding genes are non-essential in a CRISPRko screen are potential Epromoter-like elements regulating distal genes. **B**. Experimental workflow. Cells were transduced with pooled lentiviral sgRNA libraries targeting either human promoters (CRISPRi) or genes (CRISPRko), selected with puromycin, and sgRNA abundance was quantified by sequencing at Day 0 and Day 21. Genes and promoters with a negative log_2_(FC) and below a significant FDR threshold were considered CRISPRi and CRISPRko hits. **C.** Volcano plots of the CRISPRi and CRIPSRko screens. Dashed lines indicate the significance thresholds (FDR = 1%, −log_10_(FDR) = 2) and log_2_(FC) = −1. **D**. Pie chart showing the proportion of CRISPRi hits (essential promoters) that overlap or not CRISPRko hits (essential genes).

### Epromoter-like elements are enriched in promoter-promoter interactions

To study the possibility that Epromoters engage in 3D chromatin interactions, we leveraged published H3K27ac HiChIP data from Jurkat and HPB-ALL T-ALL cell lines [42]. Promoter anchors were defined by overlapping loop anchor coordinates with 1-kb windows around RefSeq transcription start sites (TSSs) (Additional file 1: Tables S2 and S3). Anchors without a TSS overlap were considered enhancers. P-P interactions accounted for 26-27% of all loops, consistent with P-P proportions found in other 3D studies [13,23,35] (Additional file 2: Fig. S1A). Notably, individual promoters interacted on average more frequently with other promoters than with enhancers, suggesting the existence of dense interacting promoter networks (Additional file 2: Fig. S2B). To further characterize the promoter-interacting networks, we clustered Jurkat interactions using the Markov algorithm (MCL), a graph-based method that simulates stochastic flow on an interaction network (Fig. 2A; Additional file 1: Table S4) [43]. Based on the number of promoters and enhancers per cluster, we obtained 1644 Enhancer-rich (E-rich), 1267 promoter-rich (P-rich) and 458 E=P clusters. Next, we checked whether the obtained cluster distribution was significantly different from random expectation by performing 1000 degree-preserving permutations of the Jurkat HiChIP interaction network and applied MCL for each one of them. We observed that the fraction of P-rich clusters (37.6%) was significantly higher than expected (33.5%, empirical *P*-value = 0.002), while the fraction of E-rich was not significantly different (Additional file 2: Fig S1C). These results reinforce the hypothesis that promoter nodes are not scattered arbitrarily, and rather tend to cluster into promoter-dominated communities. MCL produced very highly connected clusters, while our goal was to study more local interaction environments. Therefore, we adopted a promoter-centered definition for subsequent analyses, in which one promoter anchor and all its interaction partners define one promoter hub (Additional file 1: Table S5). Counting promoter and enhancer partners for each anchor yielded 4030 P-rich and 2955 E-rich hubs, again highlighting the prevalence of P-P interaction networks (Fig. 2B).

**Figure 2:**
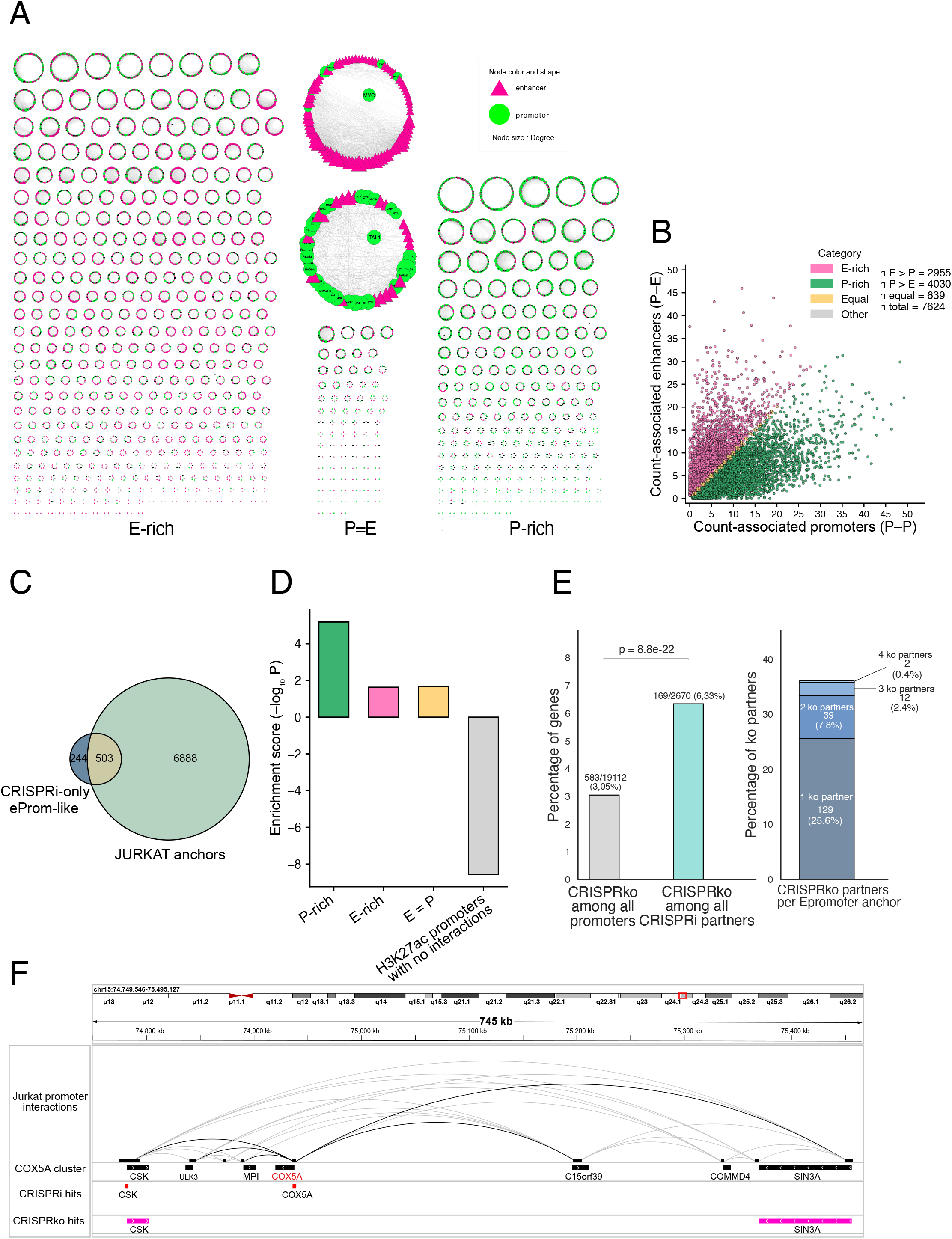
Epromoter-like elements preferentially engage in promoter-rich networks in Jurkat H3K27ac HiChIP. **A**. Markov Clustering algorithm (MCL) of the Jurkat H3K27ac HiChIP network. Clusters were classified according to the number of promoters (green circles) or enhancers (pink triangles) they contain as promoter-rich (P-rich), enhancer-rich (E-rich) or containing equal amount of both (P=E). The E-rich containing *MYC* and the P-rich containing *TAL1* are displayed enlarged. **B.** Distribution of the number of promoters and enhancers per H3K27ac-marked HiChIP anchor promoter. Anchors were classified into E-rich, P-rich, or E=P clusters. **C**. Venn diagram showing the overlap between Epromoter-like elements and Jurkat HiCHIP anchors, identifying Epromoter-like elements engaging in 3D chromatin interactions. **D**. Enrichment of the 503 Epromoter- like elements in P-rich, E-rich and E=P clusters. Epromoter-like elements are significantly enriched in P-rich clusters. *P* value was calculated using a one-sided hypergeometric test. **E**. Left, proportion of CRISPRko genes among all promoters and among all CRISPRi partners. *P* value was calculated with a one-sided hypergeometric test. Right, stacked bar plot showing the proportion of Epromoter-like hubs containing 1, 2, 3 or 4 CRISPRko partner genes. Overall, 36.18% of Epromoter-like hubs contain at least one CRISPRko anchor. **F.** HiChIP interaction map of the *COX5A* promoter cluster with tracks for CRISPRi and CRISPRko hits. Black arcs indicate interactions involving the *COX5A* hub, whereas grey arcs represent the remaining promoter-promoter interactions within the cluster.

We then intersected Jurkat promoter anchors with our CRISPRi Epromoter-like set and identified 503 Epromoter-like anchors engaging in 3D interactions (Fig. 2C). These Epromoter-like anchors were preferentially enriched in P-rich, rather than E-rich or E=P clusters (Fig. 2D), supporting the idea that Epromoter-like elements act within promoter networks and may modulate the expression of distal essential genes. To assess this hypothesis, we computed the percentage of interacting partners that are CRISPRko hits. Indeed, we found a significant enrichment for CRISPRko hits among all CRISPRi partners (Fig. 2E, left panel). For example, the *COX5A* cluster contains two essential genes (Fig. 2F), including *SIN3A,* which is a chromatin regulator known to be essential for T cell survival [44]. Overall, 36% of Epromoter-like hubs interact with at least one CRISPRko partner gene (Fig. 2E, right panel). The remaining cases might be explained by the additive effect of interacting genes.

### Promoter interactions and hubs show co-regulation in T-ALL cells

To first assess whether promoter-promoter interactions impact on gene expression, we considered unique anchors that establish specific clusters in Jurkat versus HPB-ALL cell lines (Fig. 3A). We assessed differential gene expression by analyzing RNA-seq from both cell lines [45] and found that log2 fold change was significantly greater in Jurkat for specific anchors and their cluster partners (Fig. 3B). We also observed a significant increase when we drilled down for CRISPRi-only Jurkat-specific anchors and their partners (Fig. 3B). These results indicate that emergence of new 3D interactions in one cell line leads to a specific up-regulation in gene expression.

**Figure 3.**
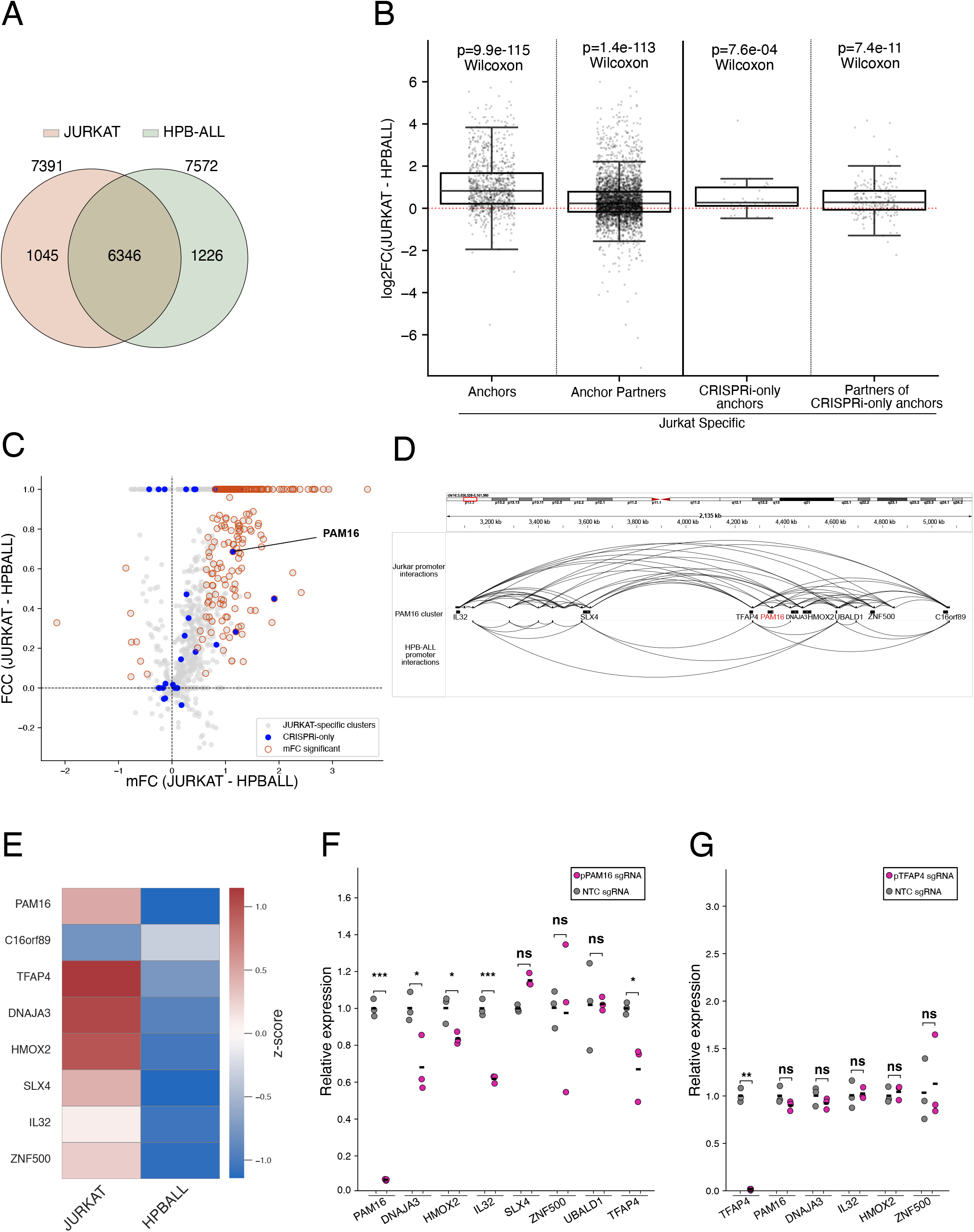
Co-regulation in the Jurkat cell-line. **A.** Venn diagram showing the number of Jurkat-specific anchors relative to HPB-ALL. **B.** Log 2FC expression between Jurkat and HPB-ALL of Jurkat-specific anchors, their partners, the CRISPRi-only Jurkat-specific anchors and their partners. Per-sample Wilcoxon signed-rank test vs 0 is shown. **C.** Scatter plot of cluster Fold Change Concordance (FCC, see methods) versus mean fold change (mFC) for all Jurkat clusters (grey). Filled magenta circles are the 503 CRISPRi-only (Epromoter-like) clusters and highlighted are mFC significant after empirical test with α=0.05. **D**. Track of Jurkat specific anchor *PAM16* showing a specific promoter cluster between Jurkat and HPB-ALL. **E**. Heatmap showing z-scored expression of the genes in the Jurkat *PAM16* cluster in Jurkat and HPB-ALL cell lines. **F.** *PAM16* CRISPRi inhibition and relative expression of the *PAM16* target genes. Dots show individual qPCR measurements and horizontal bars indicate means; significance was assessed using a two-sided Welch’s test on ΔΔCt values comparing each sgRNA sample with non-targeting control (NTC) (*p<0.05, **p<0.01, ***p<0.001).

We have previously shown that Epromoters coregulate stress-response gene clusters [32,33]. We wondered whether the genes inside the HiChIP Jurkat Epromoter clusters could also be co-regulated (Fig. 3C). Gene co-regulation has been described inside active topologically associated domains (TADs) in human cancers [46]. To address co-regulation in Jurkat clusters, we used the same strategy as in Zufferey et al. [46], computing a fold-change concordance score (FCC) for all genes in a hub that quantifies the agreement in both the sign and magnitude of the hub’s gene expression fold changes in a cell-line-specific manner. The FCC score is equal to 1 when all genes show full concordance and approaches 0 when genes show no concordance patterns. We also computed the mean Fold Change (mFC) between Jurkat and HPB-ALL expression for the genes within each Jurkat cluster and assessed statistical significance using an empirical permutation test (Fig. 3C; Additional file 1: Table S6). Overall, FCC distributions were skewed toward positive mFC, suggesting that co-regulation preferentially occurs for gene activation. Among all hubs, 302 (4%) displayed FCC>0.5 and significant mFC>1, including 32 Epromoter-like clusters (Fig. 3C). We further focused on the hub mediated by the *PAM16* Jurkat-specific anchor anchor (FCC= 0.68 and mFC=1.13) and observed higher expression of all genes in the cluster in Jurkat compared to HPB-ALL (Fig. 3D and 3E). Targeting the *PAM16* promoter with an sgRNA in Jurkat dCas9-KB cells led to extinction of PAM16 expression and significant down-regulation of *DNAJA3*, *HMOX2*, *IL32* and *TFAP4* (Fig. 3F). However, targeting the *TFAP4* promoter did not affect *PAM16* expression nor the rest of its partners (Fig. 3G). These results support that *PAM16* promoter functions as an Epromoter that regulates 3D interaction targets and contributes to hub-level co-regulation.

### Promoter hubs co-expressed in a cohort of T-ALL patients are functionally regulated

To further explore the physiological relevance of promoter hubs, we assessed the co-regulation of the interactions defined in Jurkat in a cohort of 72 T-ALL samples for whom we obtained H3K27ac and H3K4me3 ChIPseq as well as RNA-seq data (160 RNA-seq samples [47]. Widespread, positively correlated P-P interactions for active epigenetic marks have been described across diverse cell/tissue types [36]. We therefore examined whether this phenomenon generalizes to the patient cohort by computing Pearson correlations across patients for each P-P pair. Compared to random P-P pairs, interacting promoters showed significantly stronger correlation for both active chromatin marks (H3K27ac and H3K4me3) and gene expression (Fig. 4A). As an example, promoters KBN1 and CFAP20 display high correlation of the H3K27ac mark among all patients (Additional file 3: Fig. S2A). These correlations were significantly higher for P-P pairs belonging to P-rich hubs than to E-rich clusters (Additional file 2: Fig. S2B), indicating that promoter-dense networks tend to behave as coordinated regulatory neighborhoods. To probe coordination at the cluster level, we compared the Jurkat cluster mean P-P correlations to size-matched randomized clusters sampled from the global P-P pool, and observed a significant enrichment for the RNA-seq correlations (Additional file 2: Fig. S2C; Fig. 4B). Although individual P-P pairs showed positive correlations for both histone marks and RNA, only RNA displayed a significant enrichment at the cluster level, suggesting that promoter hubs primarily organize transcriptional co-regulation whereas local histone marks may reflect more gene-specific tuning.

**Figure 4.**
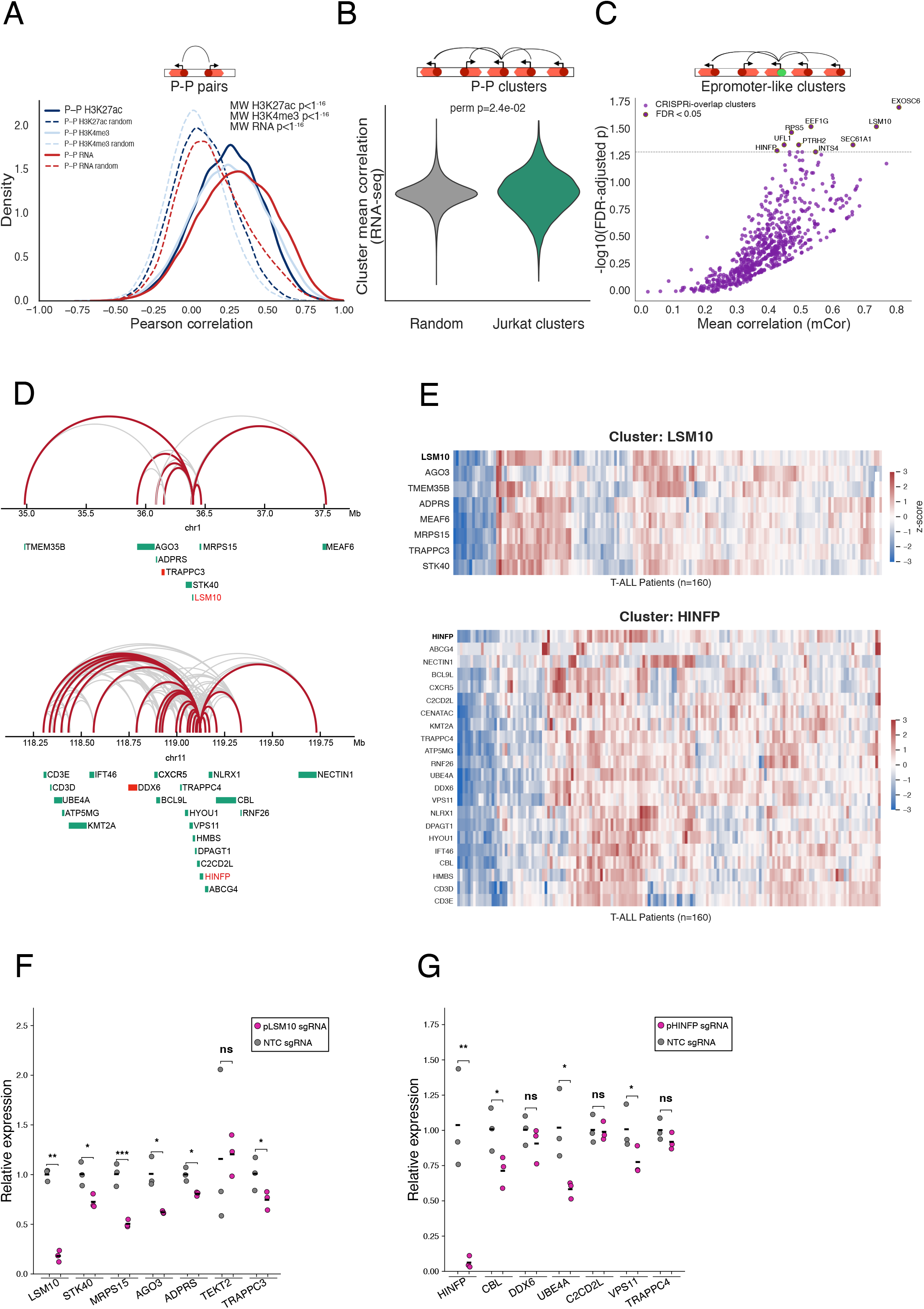
Cluster co-regulation in the T-ALL patient cohort. **A.** Kernel density plot showing the distribution of Pearson correlation coefficients for H3K27ac, H3K4me3 and RNA between individual promoter-promoter (P-P) pairs in the patient cohort. Significance was assessed using two-sided Mann-Whitney U test. **B.** Violin plot showing the distribution of cluster mean RNA correlations for Jurkat clusters and size-matched random hubs. Empirical one-sided permutation *p*-value (perm p) compares Jurkat versus random cluster mean correlations. **C.** Scatter plot of cluster means RNA correlation (mCor) versus −log_10_ (FDR-adjusted permutation p-value) for Jurkat clusters overlapping CRISPRi hits. Each point represents one cluster; highlighted dots correspond to candidate clusters with FDR <0.05 with labels for the Epromoter-like anchor. **D.** Tracks of candidate cluster interactions (red for candidate interactions, grey for all cluster interactions). Red blocks indicate a CRISPRko hit in the hub. **E**. RNA-seq expression hierarchically clustered heatmaps of the cluster genes in the 160 patients from the cohort. **F and G.** *LSM10* and *HINFP* CRISPRi inhibition and relative expression of their target genes. Dots show individual qPCR measurements and horizontal bars indicate means; significance was assessed using a two-sided Welch’s test on ΔΔCt values comparing each sgRNA sample with non-targeting control (NTC) (*p<0.05, **p<0.01, ***p<0.001).

To directly quantify cluster transcriptional co-regulation, we computed for each cluster the mean pairwise Pearson correlation (mCor) of gene expression among all the cluster genes in the cohort for 160 T-ALL RNA-seq samples. Statistical significance was assessed by performing 10,000 quantile-stratified permutations of size-matched gene-cluster assignments, thereby preserving cluster sizes and gene expression distributions, and deriving an empirical null mCor distribution per cluster with Benjamini-Hochberg FDR correction We applied this framework to all Jurkat clusters (Additional file 1: Table S7) and focused on the 503 Epromoter-like clusters (Fig. 4C). We identified 9 significantly co-regulated clusters. We expanded our analysis for two co-regulated hubs mediated by the *LSM10* and *HINFP* promoter anchors. The *LSM10* hub is located on chromosome 1 and comprises 7 genes, including *TRAPPC3*, a CRISPRko hit (Fig. 4D). The *HINFP* cluster resides on chromosome 11 and is composed of 21 genes, among which *DDX6* is a CRISPRko hit, and *KMT2A* is a known T-ALL driver (Fig. 4D). To explore the co-regulation of these candidates’ cluster, we generated hierarchically clustered heatmaps of the Epromoter and its partners in the 160 T-ALL RNA-seq cohort and identified clear coordinated blocks of expression inside patient sub-groups and for most of the genes in the cluster (Fig. 4E). Together, these results indicate that Epromoter hubs act as transcriptional co-regulation modules with coherent expression blocks across patients.

To evaluate whether the selected Epromoter candidates play a role in the co-regulation of their hubs, we experimentally inhibited them by CRISPRi targeting in Jurkat-dCas9-KB expressing cells. The RT-qPCR showed that inhibition of the *LSM10* promoter resulted in suppression of the *LSM10* gene as well as significant downregulation of *STK40*, *MRPS15*, *AGO3*, *ADPRS* and *TRAPPC3* interacting partners (Fig. 4F). Targeting of the *HINFP* promoter also resulted in the extinction of the *HINFP* gene and significant reduction of at least *CBL*, *UBE4A* and *VPS11* interacting partners (Fig. 4G). These results confirm that co-expressed gene clusters in a cohort of T-ALL patients are functionally regulated by Epromoter hubs.

### Candidate Epromoters are essential for T-ALL survival

Finally, we assessed whether the Epromoter candidates have an essential phenotype for T-ALL cells by performing a cell competition assay. Briefly, we mixed the same amount of Jurkat dCas9-KB cells transduced with the Epromoter-targeting sgRNA fused to a blue fluorescent protein (BFP) with cells carrying a non-targeting control (NTC) fused to mCherry. We used flow cytometry at days 0 to 21 to evaluate BFP/mCherry ratios (Fig. 5A). We found a drastic diminution of the amount of *LSM10*, *HINFP* and *PAM16* promoter-targeted cells from day 7 and an almost complete disappearance at day 21 (Fig. 5B-C; Additional file 2: Fig. S3). To better understand the strong impact on cell survival observed after inhibition of these promoters, we explored the CRISPRko essentiality of the interacting partners (Fig. 5D). For the *LSM10* and *HINFP* hubs, we identified at least one CRISPRko hit (*TRAPPC3* and *DDX3*, respectively), but we also found several genes with significant FDRs although they did not pass the negative fold change threshold. However, these genes could also additively contribute to the survival phenotype. For the *PAM16* hub, we did not find any essential gene, suggesting an additive or synergistic interaction between the partner genes. Overall, we identify essential Epromoter hubs driving coregulated expression of gene clusters in T-ALL primary blasts.

**Figure 5.**
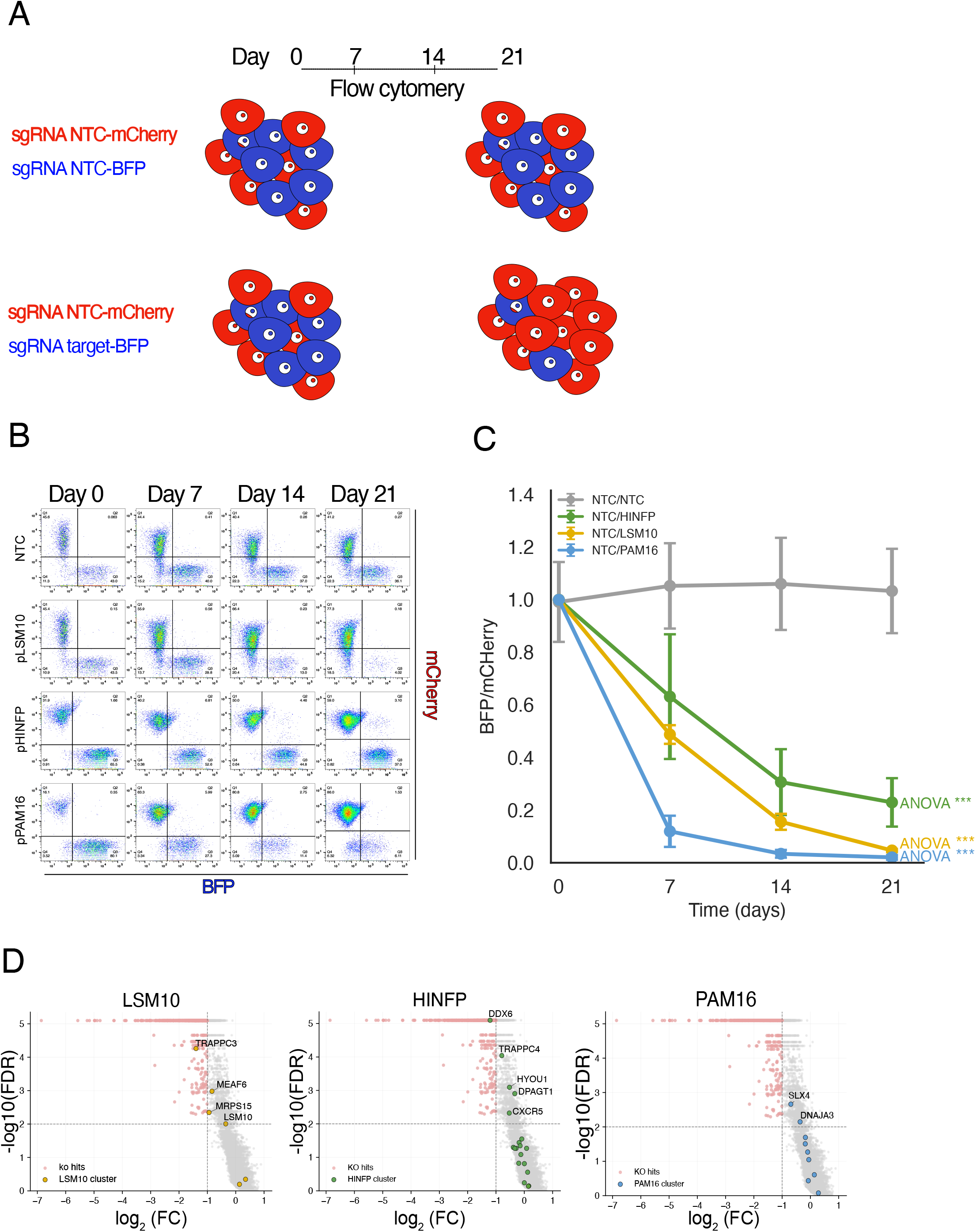
Candidate Epromoters control their targets and are essential for T-ALL survival. **A.** Schematics of competition assay. Equal numbers of transfected cells carrying the NTC sgRNA fused to mCherry and BFP, or a candidate sgRNA fused to BFP together with an NTC sgRNA-mCherry control, are mixed on day 0 and analyzed by flow cytometry on days 0, 7, 14 and 21. A negative phenotype is reflected by loss of proliferation (and progressive depletion) of cells expressing the candidate target sgRNA. **B.** Representative flow-cytometry competition assay showing the abundance of mCherry/BFP expressing cells over time. Loss of the BFP population for candidate sgRNAs (*LSM10*, *HINFP*, *PAM16*) is shown compared to NTC-transfected cells. **C.** Quantification of competition assays with ratio BFP to mCherry over time for Jurkat cells expressing candidate sgRNAs (HINFP-BFP/NTC-mCherry, LSM10-BFP/NTC-mCHerry, PAM16-BFP/NTC-mCHerry and NTC-BFP/NTC-mCherry control). Significance was evaluated by one-way ANOVA across time points for each sgRNA (***p<0.01). **D**. Volcano plots of the CRIPSRko hits. Highlighted dots are the members of each Epromoter hub. Dashed lines indicate the significance thresholds (FDR =1%, −log_10_(FDR)=2) and log_2_FC=-1

## Discussion

In this study, we identified a subset of essential Epromoters active in T-acute lymphoblastic leukemia that link cell survival with 3D gene network architecture and transcriptional coordination. We provide evidence that promoter-promoter interactions are an important layer for transcriptomic regulation of gene networks that are essential for the survival of cancer cells.

A distinctive aspect of our study is the use of parallel survival CRISPRi and CRISPRko screens to prioritize promoters that are likely to regulate distal genes. Previous strategies to identify Epromoters have relied on STARR-seq [26], eQTL analyses [36] or target-mediated CRISPR screens [27], which infer enhancer-like promoter activity from sequence or association rather than from direct survival phenotypes. In contrast, our approach uses a functional survival readout to enrich for promoters whose essentiality arises specifically from their control of distal targets rather than from local cis-effects alone. Using this strategy, we identified 747 Epromoter-like elements, representing approximately 4% of all human promoters in the CRISPRi library, consistent with previous estimates of Epromoter prevalence [34]. Thus, we deliver a transferable strategy for mapping Epromoters, and we illustrate its utility by integrating with H3K27ac HiChIP. Our findings reveal that Epromoters directly regulate their target genes through 3D physical interactions, thereby contributing to the coordinated co-regulation of promoter-centered gene clusters. Building on previous work showing that Epromoters possess enhancer activity and recruit key transcription factors to control distal, stress-responsive gene clusters [26,32,33], we now provide further evidence that these effects can be mediated via promoter–promoter (P–P) contacts. Hence, we established that Epromoter anchors were preferentially embedded in promoter-rich clusters and thus enriched in P-P interactions. These clusters are associated with increased gene expression, and the selected candidate Epromoters directly influence the expression of their interacting partners. In some cases, Epromoter essentiality could be explained by the essentiality of one or more controlled targets, as observed for *HINFP* and *LSM10* hubs and their CRISPRko target hits. In other cases, such as the *PAM16* hub, our data suggest that multiple partners may act additively or synergistically to confer essentiality. Notably, we also observed anchor directionality, exemplified by the control of *TFAP4* by the *PAM16* promoter, while inactivation of the *TFAP4* promoter did not affect *PAM16* expression, indicating that Epromoter-centered hubs encode asymmetric regulatory relationships within promoter networks.

Promoter-promoter interactions have often been proposed to primarily involve housekeeping genes [19,48,49]. In our study, we find that P-P pairs are co-regulated in T-ALL samples at both epigenetic and transcriptional levels. Moreover, the Epromoter-centered hubs that we define, comprising an Epromoter and all of its 3D interaction partners, are also highly co-regulated. These observations indicate that Epromoters drive coordinated expression of entire clusters in T-ALL. Within such hubs, Epromoters act as central nodes whose activity drives the interacting partners toward concerted transcriptional changes, consistent with prior proposals that cluster-level coregulation of genes is a key organizing principle of genome regulation [50,51]. Thus, dynamic P-P interactions mediated through Epromoters suggest that promoter-rich clusters may be essential not only for basal cellular functions but also for developmental and stress-responsive programs. In this context, active transcription hubs containing multiple genes and few enhancers have been proposed as a general principle of spatial organization [50] and several studies have shown that promoter-promoter interactions can contribute to cell-type-specific functional expression and eQTL effects [26,35,36,52]. Of note, our analysis does not attempt to deconvolute all components of these hubs, which also include enhancers, transcription factors and noncoding RNAs that together assemble higher-order regulatory complexes. Collectively, these findings argue that Epromoter-centered hubs are a versatile regulatory 3D architecture coordinating P-P interactions that coordinate gene expression beyond canonical housekeeping functions.

One key question raised by our study is whether Epromoters mediate oncogenic deregulation in cancer. Genetic alterations of Epromoters have been previously implicated in cancer [29]. For instance, a prostate cancer risk SNP have been shown to increase the enhancer properties of a promoter and deregulate distal genes with oncogenic consequences [53,54]. In our study, the candidate Epromoters anchors display functional phenotypes in T-ALL cells. One plausible mechanism is that promoter-centered 3D clusters engage oncogenic or leukemia-associated genes, such that Epromoter activity contributes to their overexpression. For example, in the *HINFP* hub, *KMT2A* represents a well-established T-ALL driver [55]. In the *PAM16* hub*, TFAP4* was already characterized as a transcriptional regulator of Galectin-9, a key mediator of the immunosuppressive tumor microenvironment in T-ALL, and emerges as a particularly compelling candidate effector [42]. An additional possibility is that Epromoter clusters define clinically relevant T-ALL subtypes. The clear co-regulation of all genes within individuals across blocks of patients suggests that hub-level expression patterns could serve as biomarkers for molecular subgroups. Taken together, these observations indicate that Epromoter-centered hubs may both drive leukemogenic gene expression programs and provide framework for subtype classifications and biomarker discovery in T-ALL.

## Conclusions

Our analysis provides support for the role of Epromoters in the regulation of 3D P-P interactions and co-regulation of promoter hubs, and suggests these interactions might play a critical role in T-ALL biology.

## Methods

### Cell culture

T-ALL Jurkat cells (ATCC) were cultured in RPMI 1640 medium (Gibco #21875) supplemented with 10% fetal bovine serum (FBS; Atlanta Biologicals). HEK293T cells were cultured in plates with DMEM high glucose (Gibco, #41,965-062) supplemented with 10% FBS. All cell lines were cultured at 37 °C in a humidified atmosphere containing 5% CO_2_.

### CRISPR screenings

Genome-wide pooled CRISPR knockout (CRISPRko) loss-of-function and CRISPR interference (CRISPRi) screens were performed using the human Brunello sgRNA library and the genome-wide CRISPRi/a v2 sgRNA library, respectively. The human Brunello CRISPR knockout library (Addgene #73179, [56]), consisting of approximately 77,000 sgRNAs targeting ∼19,000 protein-coding genes (four sgRNAs per gene), and the human CRISPRi/a v2 library (Addgene #83978, [57]), containing approximately 104,535 sgRNAs targeting ∼19,000 protein-coding genes (five sgRNAs per gene), were used for pooled screening. Target cells stably expressing Cas9 (JX17, ATCC) or dCas9-KRAB-MeCP2 (generated in-house, [58]) were transduced with pooled lentiviral sgRNA libraries at a multiplicity of infection (MOI) of approximately 0.3 to favor single sgRNA integration per cell. After Puromycin selection, cells were expanded and maintained at 50 M cells to preserve library representation of ∼500-fold coverage. Cells were collected at baseline (day 0) and after 21 days. Three independent biological replicates were performed for each experimental condition. Genomic DNA was extracted as previously described [59] and integrated sgRNA sequences were amplified using a two-step PCR strategy with primers targeting regions flanking the sgRNA cassette (Additional file 1: Table S7). The first PCR amplification was performed using KAPA HiFi DNA polymerase (Roche) to amplify integrated sgRNA sequences while preserving library complexity. For the Brunello library, the first PCR was performed using 25 independent PCR reactions, followed by a second PCR amplification using four independent reactions. For the CRISPRi/a v2 library, the first PCR was performed using 32 independent PCR reactions, followed by a second PCR amplification using four independent reactions. PCR products were purified using AMPure XP beads (Beckman Coulter). For the Brunello library, 18 PCR cycles were used for both the first and second PCR amplification steps. For the CRISPRi/a v2 library, 14 PCR cycles were used for both PCR steps. Sequencing libraries were sequenced on an Illumina NextSeq 500 TGML platform. Sequencing reads were aligned to the corresponding Brunello and CRISPRi/a v2 reference libraries, and sgRNA abundance was quantified. Differential sgRNA representation between experimental conditions was analyzed using MAGeCK ([60]) to identify genes associated with significant sgRNA enrichment or depletion. Negative hits were selected by applying a threshold of FDR < 0,001 and a negative fold change lower than 2.

### CRISPRi and CRISPRko targeting

The sgRNAs with the best fold change in the CRISPRi screen were chosen for inhibition (Additional file 1: Table S8). Oligos containing sense 5’CACC(G)+sgRNA and antisense 5’AAAC+sgRNA+(G) were synthesized, a (G) was added only if the sgRNA did not contain a G as the first nucleotide. Annealed sgRNAs were cloned into pKLV2-mCherry or Blue Fluorescent Protein (BFP) plasmids (Addgene #67977 and #67974 [61]) using BbsI-HH restriction enzyme. Lentivirus were produced by transfecting HEK293T cells with the pKLV2 vectors along with gag-pol packaging plasmid (Addgene #12260) and VSV-G envelope plasmid (Addgene #12259), using Opti-MEM (gibco, #31985-070) and TRansIT-LT1 transfection reagent (Mirus Bio, #MIR2300), and supernatant was collected at 24h and 48h. Jurkat cells stably expressing the dCas9-KRAB-MecP2 (Jurkat-dCas9-KM; [58]) or the Cas9 (JX17, ATCC) cassettes were transduced with target sgRNAs or non-targeting control (NTC) lentivirus and three days after, puromycin (2.5 µg/mL) was added for 6 days.

### Cell Competition – Flow cytometry

On day 0, non-infected, pKLV2_sgRNA-target_mCHerry/BFP and pKLV2_sgRNANTC_BFP dCas9-KB cells were counted and mixed on a ratio 1:1 (0,4M cells). 500 µL of each mix was collected, washed and resuspended in PBS. Cells were analyzed on Day 0, Day 7, Day 14 and Day 21 using a BD LSR Fortessa X-20 cytometer equipped with UV, violet, blue, yellow and red lasers. Fluorescence minus one (FMO) controls were used to define gating strategies. Each group was tested in three independent replicates and the mean ± standard deviation of the ratio BFP/mCherry was calculated relative to Day 0 and NTC sample. For each condition, overall changes across time were assessed by one-way ANOVA on replicate values with Benjamini-Hochberg correction for multiple testing. Data analysis was performed using FlowJo software.

### Gene expression analyses

Total RNA was extracted using the RNeasy Plus Mini Kit (Qiagen) following the manufacturer’s protocol. One microgram RNA was reverse transcribed using Luna Script RT Supermix (New England Biolabs, #E3010). RT-qPCR was performed using SYBR Green Master Mix (Thermo Fisher Scientific) on a QuantStudio^TM^ using 1:10 diluted cDNA. Relative expression was calculated by the 2^ΔΔCT method, normalized to *B2M* and *GAPDH* expression. Each group was tested in three independent RNA and cDNA replicates. Primers are listed in Additional file 1, Table S9.

### HiChIP H3K27ac analysis and MCL clustering

HiChIP bedpe loop interaction files were downloaded from Wiggers et al., 2025[42]. To define promoter interactions, the anchors (intervals with HiChIP H3K27ac interactions) were overlapped using bedtools intersect (v2.31.1) with the annotated RefSeq transcripts transcription start site (TSS) database, spanning ± 500 bp from TSS with a minimum of 1bp overlap. Intervals without a RefSeq overlap were considered as enhancers. For clustering, an undirected, weighted interaction graph was constructed using NetworkX, where genomic anchors represented nodes and HiChIP loop scores defined edge weights. Self-loops were added to the adjacency matrix to preserve node identity. The Markov Cluster (MCL) algorithm (*markov_clustering* Python package) was run on the HiChIP interaction graph using loop scores as edge weights (inflation parameter = 2.5) to identify densely connected interaction clusters. For visualization, HiChIP interaction loops and MCL cluster annotations were imported into Cytoscape (version 3.10.4) [62]. Nodes represented promoters and enhancers, edges represented significant HiChIP loops. To assess the statistical significance of the obtained cluster distributions, we performed label permutation tests (*n* = 1000) while keeping the graph’s topology intact. For anchor-centered hubs, connectivity files including each anchor and all its partners were created (Additional file 1: Table S5). Epromoter-like interactions were assessed using Venn diagrams using *matplotlib_venn* package in Python.

### Co-regulation analysis

#### Cell-lines

The Fold change concordance score (FCC) was computed as in Zufferey et al. [46] but applied to genes within each hub instead of a TAD, using hub definitions from Jurkat and log2 fold-changes in RNA-seq expression between Jurkat and HPB-ALL cell lines. For each hub, we also summarized the average effect size by mean log2 fold-change (mFC). Statistical significance of mFC was assessed using an empirical permutation test in which hub genes were randomly permuted within quintiles of mean partner expression, and empirical two-sided P values were calculated as the fraction of permutations with an absolute mFC greater than or equal to the observed value (with +1 added to numerator and denominator to avoid P=0), following Zufferey et al. [46].

#### Patient Cohort

ChIP-seq data from H3K27ac and H3K4me3 were previously published [47,63]. To quantify epigenomic features in promoters, promoter windows of ± 500 bp around RefSeq TSS were intersected with H3K27ac and H3K4me3 peak files using pybedtools. Consensus peaks were defined by overlapping the features present in at least two patient samples. The final promoter-associated boundaries were defined by the union of the promoter window and the overlapping consensus peaks, constrained to a maximum window of ±10 kb around the TSS. Mean signal densities within these regions were extracted from bigwig tracks via pyBigWig. ChIP-seq and RNA-seq datasets were harmonized by averaging signals across alternative promoters to generate single-gene level vectors. Downstream analysis was restricted to a shared pool of patient samples and genes common in the ChIP-seq and RNA-seq matrices. Pairwise Pearson correlation coefficients were computed across the patient cohort for P-P interactions defined in the Jurkat HiChIP dataset. To assess statistical significance, a background control model was created where each anchor was randomly coupled with another gene from the selected genomic pool. The resulting pairwise correlations for biological and random pairs were compiled into a final sample-by-interaction matrix containing the three features.

To assess cluster correlation properties, the interactions between promoter pairs per hub were combined with their corresponding H3K27ac, H3K4me3 and RNAseq correlations. Hubs containing fewer than two P-P interactions were excluded. A global background null distribution was established via 5,000 permutation iterations where in each round, random subsets of interactions equal in size to the real hub were sampled without replacement from full P-P dataset. Observed hub mean correlations were compared to the interactions null distribution to compute empirical one-sided *p*-values for enrichment.

Mean hub correlation (mCor) was calculated for each gene hub to quantify intra-hub transcriptomic co-regulation. The observed mCor was defined as the arithmetic mean of the upper-triangular elements of the Pearson correlation matrix computed across all constituent genes in the cohort expression data. To evaluate hub significance against an expression-matched background, a high-resolution gene-level permutation test was executed using 10,000 iterations. All genes within the expression dataset were stratified into five expression quintiles. For each permutation round, gene labels were shuffled exclusively within their matching quintile. An empirical two-sided *p*-value was calculated for each hub.

*P*-values from both hub analyses were corrected for multiple testing across all hubs using Benjamini-Hochberg False discovery Rate (FDR) with a significance threshold at alpha = 0.05.

## Abbreviations

ANOVA: analysis of variance
AP-HP: Assistance Publique–Hôpitaux de Paris
BFP: blue fluorescent protein
3D: three-dimensional
cDNA: complementary DNA
ChIP-seq: chromatin immunoprecipitation followed by sequencing
CREs: cis-regulatory elements
CRISPR: clustered regularly interspaced short palindromic repeats
CRISPRi: CRISPR interference
CRISPRko: CRISPR knockout
DepMap: Dependency Map (Cancer Dependency Map)
DMEM: Dulbecco’s modified Eagle medium
E=P: enhancer = promoter (balanced clusters)
E-P: enhancer–promoter
E-rich: enhancer-rich
eQTLs: expression quantitative trait loci
FCC: fold-change concordance score
FBS: fetal bovine serum
FDR: false discovery rate
FMO: fluorescence minus one
H3K4me3: histone H3 lysine 4 trimethylation
H3K27ac: histone H3 lysine 27 acetylation
INEM: Institut Necker-Enfants Malades
Inserm: Institut national de la santé et de la recherche médicale
mCor: mean pairwise Pearson correlation
MCL: Markov Cluster (algorithm)
mFC: mean fold change
MOI: multiplicity of infection
NTC: non-targeting control
P-P: promoter–promoter
PBS: phosphate-buffered saline
P-rich: promoter-rich
RT-qPCR: reverse transcription quantitative polymerase chain reaction
sgRNA: single-guide RNA
T-ALL: T-cell acute lymphoblastic leukemia
TADs: topologically associating domains
TAGC: Theories and Approaches of Genome Complexity
TSS: transcription start site

## Declarations

### Ethics approval and consent to participate

No ethical approval or patient consent was required for this study.

### Consent for publication

Not applicable.

### Availability of data and materials

The datasets supporting the conclusions of this article are available under the number GSE344160 at the GEO repository, https://www.ncbi.nlm.nih.gov/geo/query/acc.cgi?acc=GSE344160.

The datasets supporting the conclusions of this article are included within the article and Additional file 1.

### Authors’ contributions

JFRP and SS conceptualized the study. JFRP, IM, CC and NS performed the experiments. JFRP performed bioinformatic analyses, downstream data analysis and generated figures and tables. AZ contributed to network analysis. AP, AC, GA, CA, VA and AT contributed to patient annotation. JFRP and SS wrote the manuscript with inputs from the authors.

## Acknowledgements

We thank the Marseille-Luminy cell biology platform for the management of cell culture (PCC), the Transcriptomics and Genomics Marseille-Luminy (TGML) sequencing platform and the CRISPR Screen Action platform from the Canceropôle PACA. We acknowledge the contribution of SFR Biosciences (UAR3444/CNRS, US8/Inserm, ENS de Lyon, UCBL) facilities, particularly the AniRA lentivector production facility from the CELPHEDIA Infrastructure (Gisèle Froment and Caroline Costa). We thank the members of TAGC for helpful discussions and critical reading of the manuscript.

---

**Figure S1.**
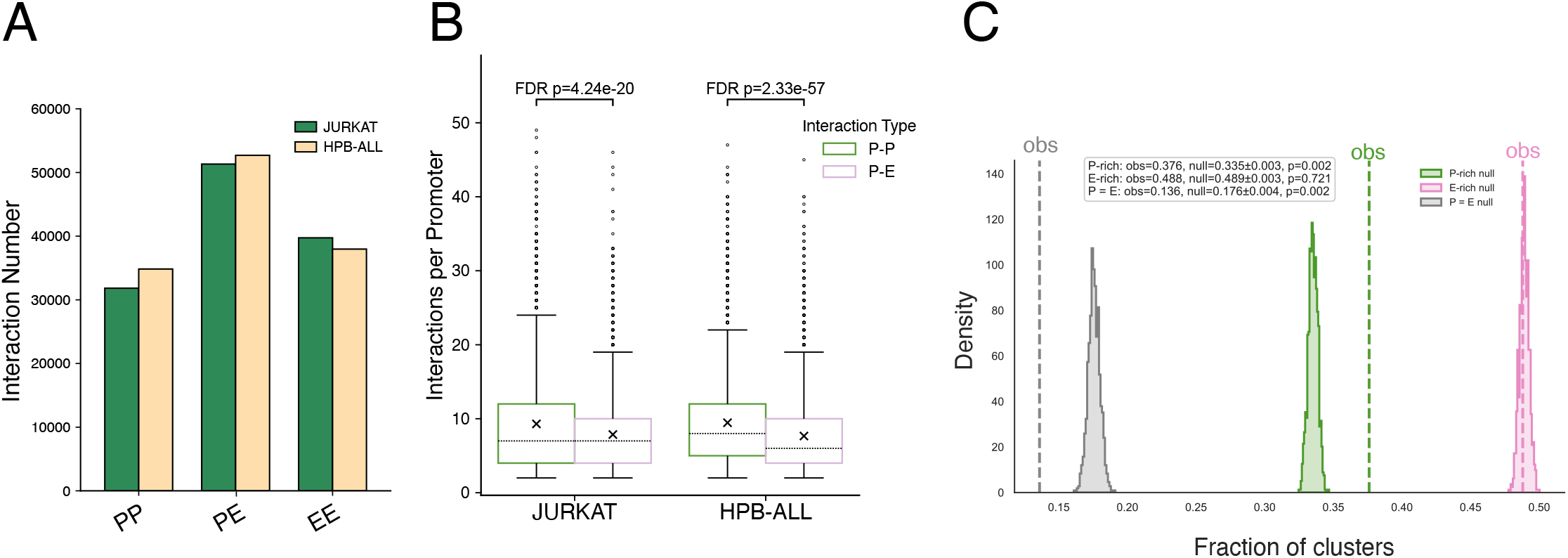
Promoter-rich interactions in Jurkat and HBPALL H3K27ac HiChIPs. **A.** Number of promoter-promoter (PP), promoter-enhancer (PE) and enhancer-enhancer interactions (EE) in Jurkat and HPB-ALL H3K27ac HiChIP networks. **B**. Average number of interactions per promoter in Jurkat and HPB-ALL cells. Crosses indicate the mean number of interactions and the horizontal dashed lines within each box indicate the median. Statistical significance within each cell-line was assessed using a Mann-Whitney U test with Benjamini-Hochberg false discovery rate (FDR) correction. **C**. Density plot showing the distribution of Markov clustering (MCL) results from 1000 degree-preserving randomizations of the Jurkat HiChIP interaction network. Filled curves represent the empirical null distributions of the fraction of MCL clusters in each category, and dashed lines indicate the corresponding observed fractions in the real network. Statistical significance was assessed using two-sided empirical *P* values derived from null distributions.

**Figure S2.**
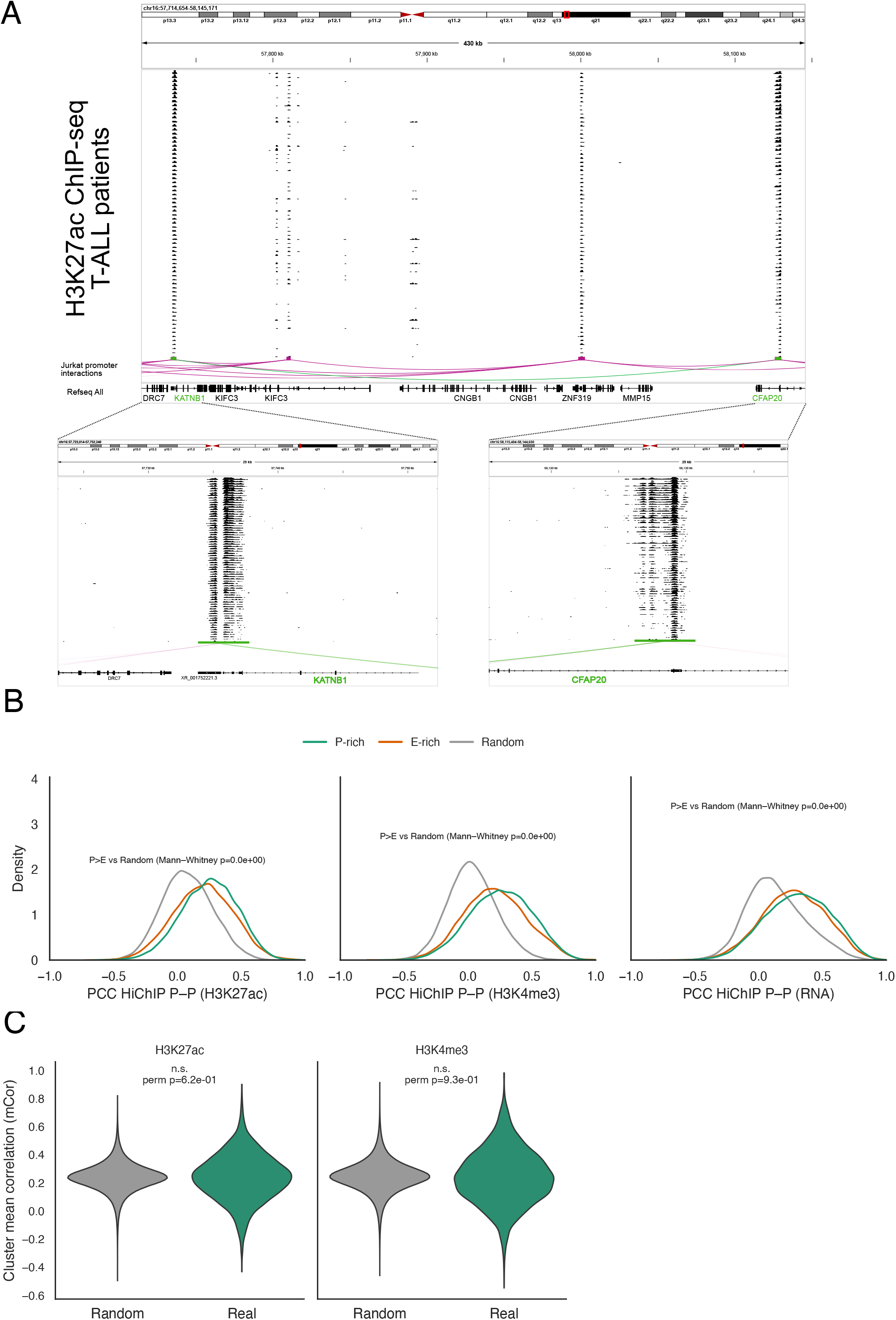
Cluster co-regulation in the T-ALL patient cohort. **A.** Example of correlated P-P pair KATNB1 and CFAP20 showing tracks for H3k27ac ChIP-seq among the patient cohort**. B.** Kernel density plots showing the distribution of Pearson correlation coefficients (PCC) for H3k27ac, H3K4me3 and RNA between P-rich (green) and E-rich (orange) vs random (grey) in the patient cohort. Significance was assessed using a two-sided Mann-Whitney U test. **C.** Violin plots showing the distribution of cluster mean H3K27ac and H3k4me3 correlations for Jurkat clusters and size-matched random hubs. Empirical one-sided permutation p-value (perm p) compares Jurkat versus random cluster mean correlations (* p<0.05).

**Figure S3.**
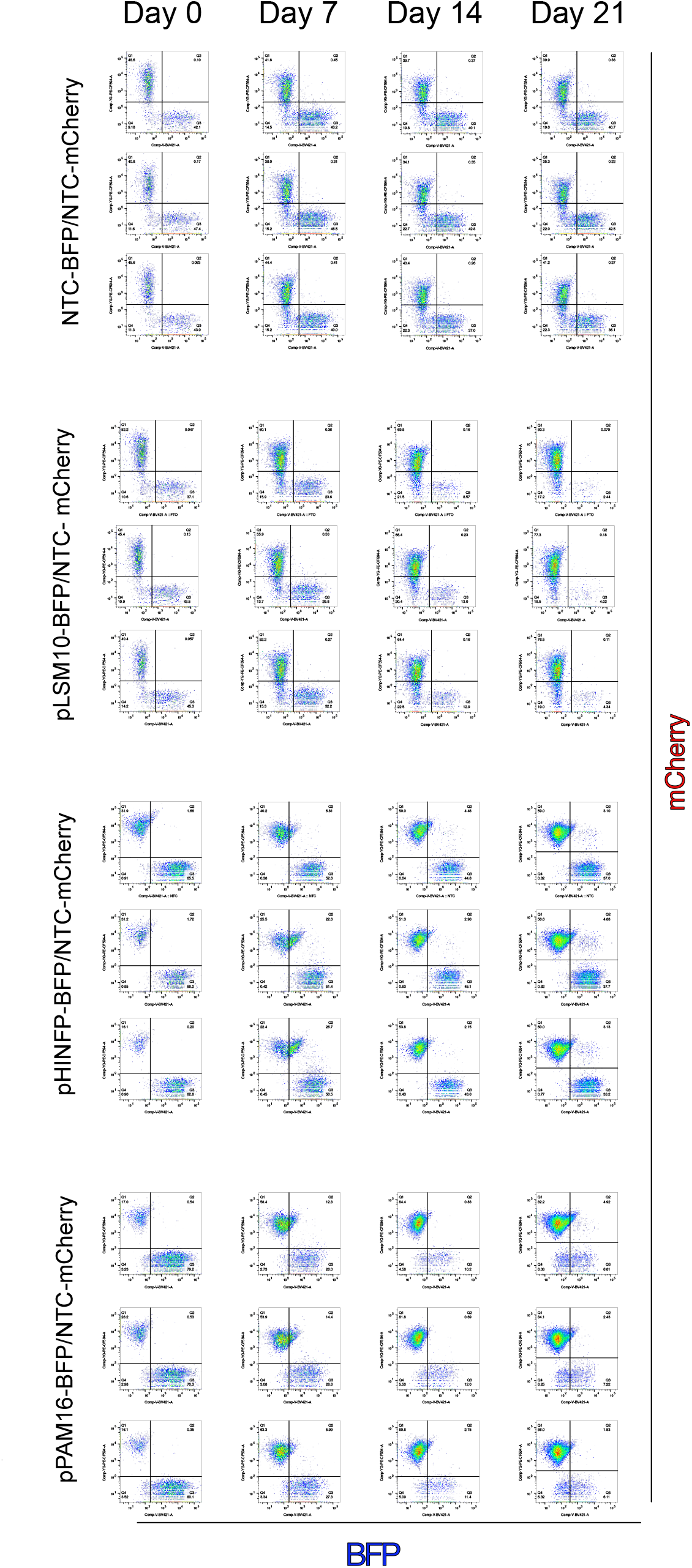
Complete set of biological replicates of flow-cytometry competition assays showing the abundance of mCherry/BFP expressing cells over time. Loss of the BFP population for candidate sgRNAs (LSM10, HINFP, PAM16) is shown compared to NTC-transfected cells.

